# Interaction of Sox2 with RNA binding proteins in mouse embryonic stem cells

**DOI:** 10.1101/560383

**Authors:** Samudyata, Paulo P. Amaral, Pär G. Engström, Samuel C. Robson, Michael L. Nielsen, Tony Kouzarides, Gonçalo Castelo-Branco

## Abstract

Sox2 is a master transcriptional regulator of embryonic development. In this study, we determined the protein interactome of Sox2 in the chromatin and nucleoplasm of mouse embryonic stem (mES) cells. Apart from canonical interactions with pluripotency-regulating transcription factors, we identified interactions with several chromatin modulators, including members of the heterochromatin protein 1 (HP1) family, suggesting a role of Sox2 in chromatin-mediated transcriptional repression. Sox2 was also found to interact with RNA binding proteins (RBPs), including proteins involved in RNA processing. RNA immunoprecipitation followed by sequencing revealed that Sox2 associates with different messenger RNAs, as well as small nucleolar RNA Snord34 and the non-coding RNA 7SK. 7SK has been shown to regulate transcription at regulatory regions, which could suggest a functional interaction with Sox2 for chromatin recruitment. Nevertheless, we found no evidence of Sox2 modulating recruitment of 7SK to chromatin when examining 7SK chromatin occupancy by Chromatin Isolation by RNA Purification (ChIRP) in Sox2 depleted mES cells. In addition, knockdown of 7SK in mES cells did not lead to any change in Sox2 occupancy at 7SK-regulated genes. Thus, our results show that Sox2 extensively interact with RBPs, and suggest that Sox2 and 7SK co-exist in a ribonucleoprotein complex whose function is not to regulate chromatin recruitment, but might rather regulate other processes in the nucleoplasm.

**Summary blurb:** Sox2 interacts with RNA-binding proteins and diverse RNAs

## Introduction

The defining features of embryonic stem (ES) cells are self-renewal and pluripotency, both of which are governed by complex gene regulatory networks. The master transcriptional regulator, Sox2 (SRY-box containing gene 2) lies at the center of these programs (Avilion et al., 2003; Takahashi and Yamanaka, 2006). Sox2 binds to DNA via its highly conserved HMG-box domain, often in co-operation with other transcription factors of the pluripotency network, such as Oct4 and Nanog (Avilion et al., 2003; Gao et al., 2012), to elicit programs that either maintain ES cell identity or lead towards differentiation of multiple lineages (Wang et al., 2012; Zhang and Cui, 2014). ES cells harbour a unique epigenetic landscape defined by permissive chromatin with a more dispersed heterochromatin along with bivalent histone marks placed on developmentally important genes (Gaspar-Maia et al., 2011). This plasticity forms a crucial part of the regulatory circuit and is contributed by a dynamic and reciprocal interaction of epigenetic modulators such as histone/DNA modifiers and nucleosome remodellers with the core pluripotency transcription factors in ES cells (Delgado-Olguín and Recillas-Targa, 2011; Guenther et al., 2010; Kashyap et al., 2009). This cross-talk between key transcription factors, such as Sox2, and chromatin modulators also occurs in other multipotent cells types, such as neural stem cells (Engelen, Akinci et al., 2011). Non-coding RNAs (ncRNAs) have also emerged as important regulators of chromatin status and transcription and are likely to operate within a highly integrated network of transcription factors and chromatin modulators to influence key cellular events (Huo and Zambidis, 2013; Wright and Ciosk, 2013).

In this study, we identified several chromatin modulators and RNA binding proteins interacting with Sox2 in different nuclear fractions of embryonic stem (ES) cells, by Stable Isotope Labelling by Aminoacids in Cell culture (SILAC) technology (Ong et al., 2002), coupled with immunoprecipitation and mass spectrometry-based quantitative proteomics. In addition, we affinity-purified Sox2 from mES cell extracts and identified associated RNAs through RNA-sequencing, including the small nuclear RNA (snRNA) 7SK and small nucleolar RNA (snoRNA) Snord34. 7SK is known to regulate transcriptional elongation by sequestering positive transcription elongation factor b (P-TEFb), a critical factor required for Pol II promoter proximal pause-release, in a catalytically inactive small nuclear ribonucleoprotein complex (Peterlin et al., 2012). We have previously shown that 7SK can regulate genes involved in lineage commitment, suggesting directed recruitment to specific regulatory regions in mES cells (Castelo-Branco et al., 2013). Nevertheless, we could find no evidence of Sox2 regulating 7SK recruitment to chromatin, or vice-versa, suggesting that the interactions between 7SK and Sox2 might be involved in other processes. In sum, our data suggests that Sox2 is present in complexes containing chromatin regulators and RNA binding proteins, which indicates that Sox2 may be involved in their functions and that its role as a transcriptional regulator might involve association with specific RNAs.

## Results

Sox2 has been shown to be a key player in maintaining the pluripotent state of ES cells. In order to identify the protein complexes associated with Sox2 in mouse pluripotent cells, we combined affinity purification of biotin-tagged recombinant proteins with SILAC quantitative proteomics (Figure 1A). To explore the protein interactors of Sox2 in different nuclear fractions, we prepared native chromatin and nucleoplasm extracts of ^13^C_6_-labelled J1 ES cells expressing Sox2 biotinylated by BirA (bioSox2) and ^12^C_6_-labelled J1 control ES cells, expressing only BirA. Protein complexes interacting with Sox2 were immunoprecipitated with streptavidin beads and mixed 1:1 with control samples prior to proteomic analysis by mass spectrometry.

**Figure 1.**
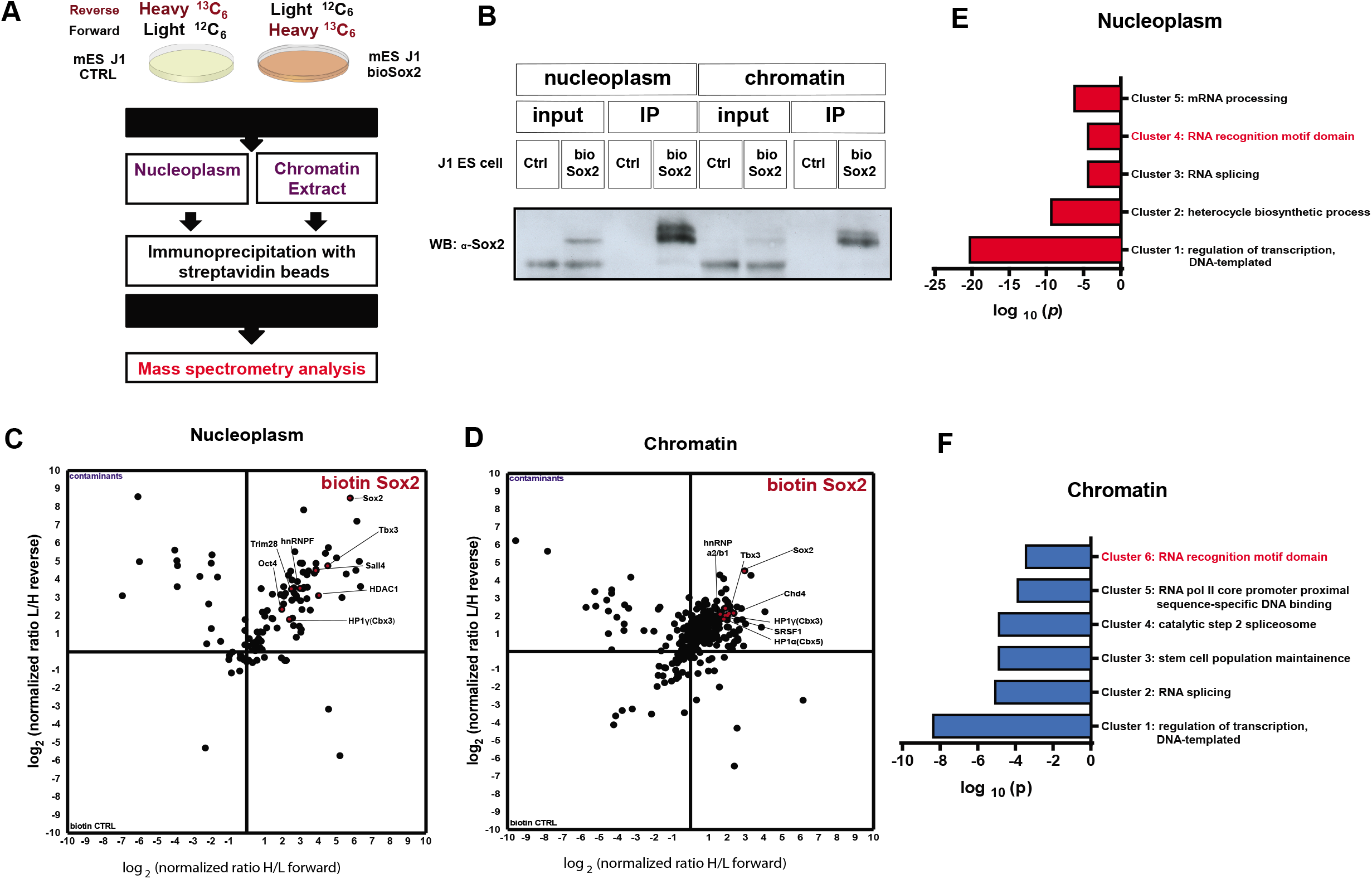
A) Schematic representation of the strategy used to characterize Sox2 protein interactome by Stable Isotope Labelling by Amino acids in Cell culture (SILAC) followed by Mass spectroscopy (MS). Control and bioSox2 J1 ES cell lines were cultured with either LIGHT (^12^C_6_) or HEAVY (^13^C_6_) medium, respectively. Native chromatin and nucleoplasm extracts were prepared from these cells and the protein interactome of Sox2 was immunoprecipitated and mixed prior to MS for proteomic analysis. B) Western blot showing successful pull down and an enrichment of bioSox2 after immunoprecipitation in both chromatin and nucleoplasm fractions, compared to the control. C) 2D interactome plot representing the fold change of identified proteins interacting with bioSox2 in the nucleoplasm. Ratios are represented in a logarithmic scale with (H/L) on X axis plotted against (L/H) on Y. D) 2D interactome plot representing the fold change of identified proteins interacting with bioSox2 in the chromatin. Ratios are represented in a logarithmic scale with (H/L) on X axis plotted against (L/H) on Y. E) Gene Ontology (GO) analysis for significant protein interactors of Sox2 in the nucleoplasm fraction of J1 ES cells. Represented in the figure are the non-redundant GO terms found over-represented by modified Fisher exact test with Bonferroni corrected P-values F) GO analysis for significant protein interactors of Sox2 in the chromatin fraction of J1 ES cells. Represented in the figure are the non-redundant GO terms found overrepresented by modified Fisher exact test with Bonferroni corrected P-values.

For increased specificity, we also performed reverse labelling (^13^C_6_-labelled J1 control ES cells and ^12^C_6_-labelled bioSox2 J1 ES cells). As previously reported (Wang et al., 2006), the levels of biotinylated Sox2 were lower than endogenous Sox2 (Figure 1B). In order to determine if the somewhat elevated Sox2 expression led to ectopic differentiation, as previously reported (Kopp et al., 2008), transcriptomic profiles of bioSox2 and control J1 mES cell lines were compared and were found to be very similar (Pearson correlation coefficient R = 0.97; Supplementary Figure 1A). Amongst the few genes that were differentially expressed between the two cell lines, there was Sox21 whose elevated expression have been previously reported to trigger ES cell differentiation (Mallanna et al., 2010). Nevertheless, bioSox2 cells exhibited an undifferentiated morphology in culture (not shown) and no other differentiation markers were found to be enriched in bioSox2 compared to its control cell line (Supplementary Table 1).

For quantitative proteomics comparisons, proteins that showed at least two-fold enrichment in bioSox2 over control in both forward and reverse labelling were considered for analysis. As expected, Pou5f1 (Oct4), one of the master transcription factors of the core pluripotency network as well as other partner factors involved in stem cell maintenance such as Tbx3, Sall4, Esrrb and members of the Klf family of transcription factors were found to interact with Sox2 in the nucleoplasm (Figure 1C, Supplementary Table 2). Tbx3 and Sall4 were also found in the chromatin fraction (Figure 1D, Supplementary Table 3). Several chromatin remodelers such as Brg1-associated factors (Baf60a, Baf155, Baf57) and Chd4 (catalytic subunit of Nucleosome-remodelling comlex (NuRD)), essential for ES cell renewal, along with other chromatin modifiers like HP1 α, β, γ (Cbx5, 1 and 3), Myst4, Sin3a, Kdm5b, Pcgf2 and Eed were recovered in the chromatin fraction (Figure 1D, Supplementary Figure 1B, Supplementary Table 2). Interestingly, we could also find Sox2 association with chromatin regulators such as Trim28, Hdac1 and HP1γ in the nucleoplasm fraction. We confirmed the interaction of Sox2 with HP1 proteins using recombinant human Sox2 (Supplementary Figure 1C) or ES cell nucleoplasm extracts (Supplementary Figure 1D). To further investigate the nature of these interactions, domains from both HP1α and HP1β along with their full lengths were used to co-immunoprecipitate recombinant Sox2. Different domains in both proteins contributed towards interacting with Sox2 (Supplementary Figure 1E).

Analysis of gene ontology terms confirmed that Sox2 interactors were enriched for regulators of transcription, but also indicated that a subset of the interactors had RNA recognition motifs (Figure 1E and F). Indeed, heterogenous nuclear riboproteins such as hnRNPM, hnRNPC1/C2, hnRNPF, hnRNP2 (Fox2), hnRNPD0, hnRNPH1, hnRNPU and other RNA binding proteins involved in splicing/post-transcriptional processes such as Ddx3, Ddx5 and Ddx17 were detected as Sox2 interactors in the nucleoplasm fraction, while Fubp2, Fubp3, Rbm38, hnRNPA2/B1, Prp19, Prp8, Magoh and Srsf1 were detected in the chromatin fraction (Supplementary Tables 2 and 3). Moreover, many of the chromatin regulators observed to interact with Sox2 have been shown to interact with RNA, including HP1 (Muchardt et al., 2002). Nevertheless, we observed that the interaction between Sox2 and HP1α/β persisted upon RNAse A treatment (Supplementary Figure 1D), indicating that the observed interaction is not dependent on RNAs. In sum, these data suggest that Sox2 can be a component of ribonucleoprotein complexes in mES cells.

To examine which RNAs could be associated with these complexes, we performed two independent immunoprecipitations of bioSox2 from formaldehyde cross-linked J1 ES cells, followed by poly(A)-neutral RNA-seq (Figure 2A). While long ncRNAs were not found enriched upon Sox2 pull down, we detected an enrichment of a restricted subset of RNAs (Figure 2E and Supplementary Table 4), including mRNAs and two non-coding RNAs, the snRNA 7SK and snoRNA Snord34 in both experiments (Figure 2B and Supplementary Table 4). In order to validate the interaction of 7SK and Snord34 RNAs with Sox2 protein, we performed qRT-PCR following RNA immunoprecipitation (RIP) with biotinylated Sox2, Oct4 and Nanog, as well as RIP with antibodies against endogenous Sox2 and other pluripotency transcription factors (Supplementary Figure 1F, G). These experiments confirmed the pulldown of 7SK and Snord34 by Sox2. We observed that immunoprecipitation of other pluripotency transcription factors, such as Oct4, Nanog and Klf4 could also pull down these non-coding RNAs, albeit to a lower extent, in line with their co-existence in complexes in the nucleus (Supplementary Figure 1F). Interestingly, we found specific interaction of transcription factors with their own mRNA (except for Sox2 mRNA) (Supplementary Figure 1G), which could be due to crosslinking of the mRNA and protein during translation, or reflect recruitment of the mRNA by the respective transcription factor, in a similar manner as it has been described in Drosophila for proteins of the male-specific lethal (MSL) complex (Johansson et al., 2011).

**Figure 2.**
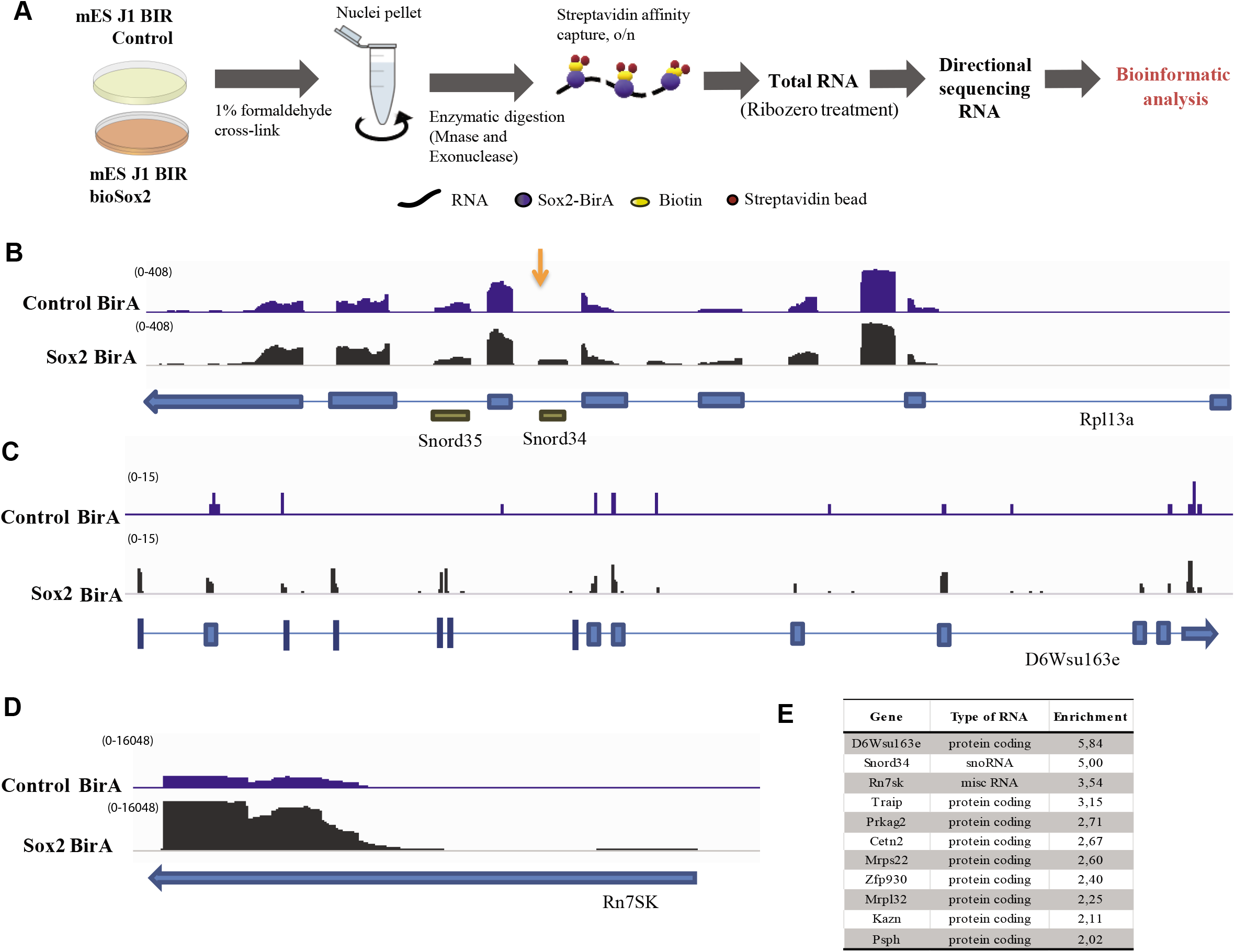
A) Schematic representation of the strategy used to characterize RNA interactome of Sox2 by RNA-immunoprecipitation followed by sequencing (RIP-seq). Cells from control and bioSox2 J1 ES cell line were fixed with 1 % formaldehyde to capture direct and indirect RNA-bioSox2 interactions. Nuclei were pelleted and RNA was enzymatically digested. BioSox2-bound RNA was immunoprecipitated with streptavidin beads and the final eluted RNA was Ribo-Zero treated to remove ribosomal RNA, before sequencing. B) IGV screenshot of *Rpl13a* gene from one RIP-Seq experiment showing normalized read counts from sequenced RNA in control and Sox2-BirA (bioSox2) samples, following RIP-seq. *Snord34* reads are over-represented in bioSox2 compared to the control (indicated by an arrow). Neighboring *Snord35* does not show any such overrepresentation. C) IGV screen shot of *D6Wsu163e* gene from one RIP-Seq experiment showing normalized read counts from sequenced RNA in control and Sox2-BirA (bioSox2) samples, following RIP-seq. *D6Wsu163e* reads are over-represented in bioSox2 compared to the control sample. D) IGV screen shot of *Rn7SK* gene from one RIP-Seq experiment showing normalized read counts from sequenced RNA in control and Sox2-BirA (bioSox2) samples, following RIP-seq. *Rn7SK* reads are over-represented in bioSox2 compared to the control sample. E) Table showing all RNAs with enrichment ratio > 2 and bioSox2 RIP raw read count > 50 in two RIP-seq experiments combined. Enrichment ratios were computed as (bioSox2 RIP / ctrl RIP) / (bioSox2 input / ctrl input), using normalized counts incremented by a pseudo-count of 0.1 (to avoid denominators of zero). For more details, see Supplementary Table 4.

Recently, 7SK was shown to occupy promoters and enhancers to regulate transcription via association with different molecular partners (Flynn et al., 2016). Given that they are both transcriptional regulators, the observed interaction between 7SK and Sox2 could play a role in their recruitment to the chromatin. To assess whether genomic recruitment of 7SK is altered in the absence of Sox2, we performed Chromatin Isolation by RNA Purification (ChIRP) with even and odd sets of probes to 7SK (Flynn et al., 2016) in a doxycycline inducible Sox2-knock out mES cell line and compared it with controls treated with DMSO. As a negative control, a single probe against LacZ mRNA was used (Figure 3A). We efficiently retrieved 7SK, although the percentage of retrieval was variable between odd and even pools (Figure 3B), as previously reported for ChIRP experiments (Chu, Qu et al., 2011). 7SK-specific probes did not retrieve GAPDH or the abundant nuclear ncRNA MALAT1, and the negative control showed negligible enrichment of 7SK ncRNA (Figure 3B). Consistent with previous reports (Chu et al., 2011), the overlap between odd and even probes in ChIRP was low. We nevertheless could identify 583 robust peaks common to both odd and even data sets but depleted for LacZ binding, in DMSO and doxycycline treated samples (Supplementary Table 5). However, we could not detect any change in the levels of 7SK binding at these common peaks following doxycycline induced Sox2 KO when compared to the control conditions (t = −0.69, df = 1.45, p = 0.585, Figure 3C). Therefore, Sox2 appears not to be involved in the recruitment of 7SK snRNA to chromatin.

**Figure 3.**
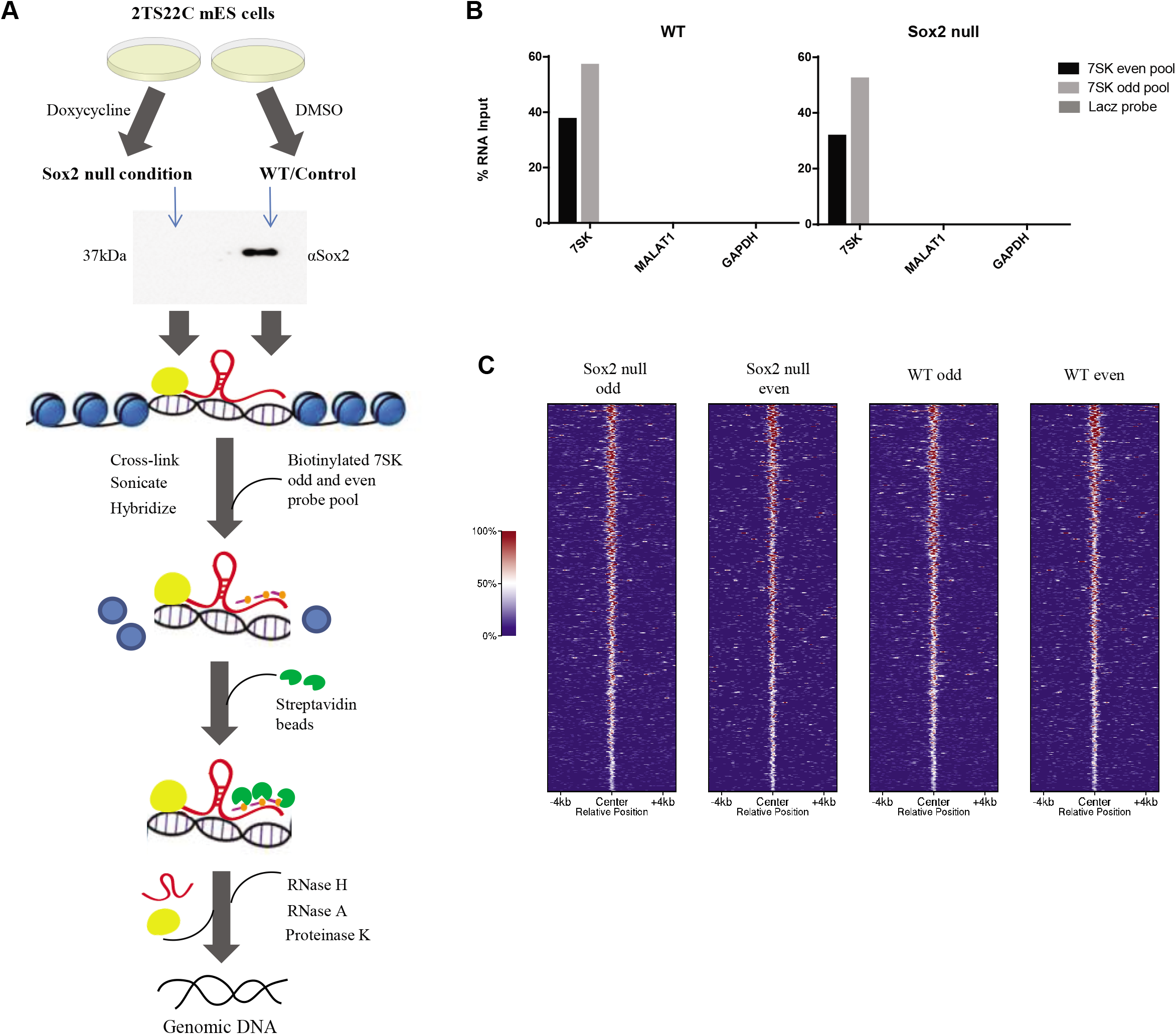
A) Schematic representation of Chromatin Isolation by RNA Purification (ChIRP) strategy to assess global recruitment of 7SK to the chromatin following Sox2 KO. Doxycycline inducible Sox2 KO 2TS22C mES cells were treated with 1 μg/ml doxycycline or DMSO for 24hrs. Western blot with 30 ng of protein extract from doxycycline treated and untreated cells shows a deletion of Sox2 in the treated samples. Sox2 null and WT 2TS22C cells were cross-linked with glutaraldehyde, sonicated and hybridized to 7SK odd and even biotinylated pools (three probes per pool) or a single biotinylated probe against LacZ mRNA. Streptavidin beads were used to pull down DNA bound by 7SK and then sequenced. B) RT-qPCR showing percent RNA pulled down following ChIRP with 7SK odd and even pools in Sox2 null and WT mES cells. *7SK* is pulled down specifically with varying efficiencies by the 7SK odd and even pool compared to the LacZ control. Neither *Gapdh* nor *Malat1* RNA show any enrichment with 7SK odd and even pools in both the conditions. C) Comparison of global genomic 7SK binding in WT and Sox2 null conditions in 2TS22C cells. Heat map showing ChIRP-seq signal, normalized to read depth, +/− 5 Kb around peak mid-points common to 7SK odd and even data sets in Sox2 null and WT samples from one ChIRP experiment. There is no significant change in global genomic 7SK recruitment following Sox2 ablation.

We then investigated whether 7SK ncRNA could instead have an impact in the association of Sox2 to specific regions on the chromatin. For this purpose, Chromatin Immunoprecipitation (ChIP) was performed with an antibody against endogenous Sox2 in mES cells where 7SK was depleted with an antisense oligonucleotide (ASO) targeting its 3’ end (Castelo-Branco et al., 2013), which was then followed by qPCR (Figure 4A,D). In order to choose suitable candidate target regions, Sox2 peaks associated with annotated genes (7,055 unique genes) from previously published ChIP-Seq experiment in mES cells (Whyte et al., 2013) and 7SK occupied regions from our ChIRP dataset with 583 robust peaks (291 unique genes) and the Flynn data set with 50,071 peaks (12,896 unique genes) were compared. There was a significant overlap of 59% and 75% of Sox2 occupied genes with ours and Flynn’s ChIRP datasets, respectively (Figure 4C). Nevertheless, when centering ChIRP reads at the Sox2 binding peaks, we could not find a clear correlation between 7 SK and Sox2 occupancy (Figure 4B). Out of the 164 genes common to all datasets (Supplementary table 6), *Kdm2b, Celf2* and *Klf12* were chosen for ChIP-qPCR analysis, along with other regions known to be occupied by Sox2 (*Pouf51* and *Nanog*) or shown to be regulated upon 7SK knock down (*Dll1*) (Castelo-Branco et al., 2013). We observed Sox2 occupancy at regulatory regions of *Pou5f1* (Oct4), *Nanog, Kdm2b, Celf2* and *Klf12*, but not at the negative control (intron of Sox10) (Figure 4E). However, knockdown of 7SK (Figure 4D) did not lead to significant changes in Sox2 binding (Figure 4E). Thus, snRNA 7SK and transcription factor Sox2, though present in the same complex, are not involved in reciprocal recruitment to these specific regions of chromatin.

**Figure 4.**
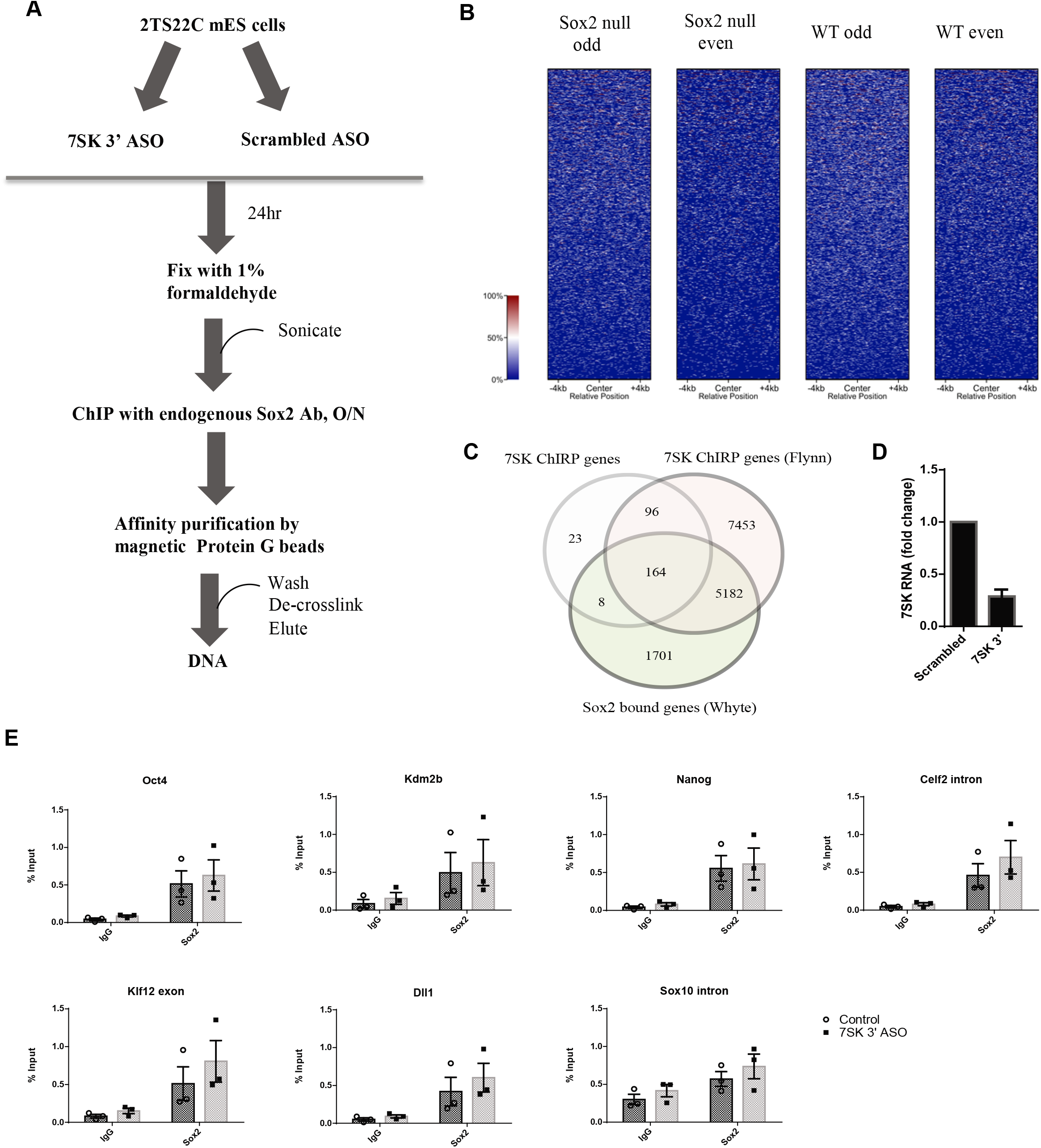
A) Schematic representation of a Chromatin immunoprecipitation (ChIP) experiment following 7SK knock down in 2TS22C mES cells. An ASO targeting 3’ end of 7SK was used to knock down 7SK. Control cells were treated with a scrambled ASO. The resulting cells were fixed with 1% formaldehyde, sonicated, and chromatin from about 100,0 cells was used for immunoprecipitation with an antibody against endogenous Sox2. This was followed by affinity purification of immune complexes with Protein G beads. The DNA was de-crosslinked and eluted prior to qPCR analysis. B) Normalized ChIRP-seq read distribution centered on the Sox2 binding peaks from Whyte *et al*. dataset shows no co-binding of 7SK at Sox2-bound loci. C) Venn diagram showing an overlap of genes among three datasets, namely Sox2 ChIP Whyte dataset, 7 SK ChIRP Flynn dataset and the ChIRP dataset produced in this study. The numbers in the intersections denote the number of unique genes associated with each factor, either in the gene body or in the promoter. D) RT-qPCR showing fold change in 7SK expression 24 h post-transfection with 100 nM of 7SK 3’ ASO compared to the control treated with a scrambled ASO. Error bars indicate SEM (n=3) E) ChIP-qPCR results showing enrichment of Sox2 bound DNA as percent input in 2TS22C cells treated with control and 7SK 3’ ASO at regulatory regions of *Pou5f1* (Oct4), *Nanog, Kdm2b, Celf2, Klf12* and *Dll1*. Amplification in goat IgG was used as a measure of background for the specific regions assayed. Sox10 intron was used as a negative control. Error bars indicate SEM (n=3), each point is a biological independent experiment (knock-down) that represents an average of triplicate or duplicate ChIP experiments.

## Discussion

Sox2 is known to exist in high molecular weight complexes, the protein interactome of which is highly dependent on the cellular context as well as on the purification and mass spectrometric methods used to isolate and determine the interactome. Our data, consisting of 124 proteins, provides a resource for the interactome of Sox2 in mESCs in different nuclear fractions. About 23% of our Sox2 interactors overlap with previously published Sox2 interactome data from the studies of Gao et al. and Mallana et al. (Supplementary Table 7). Given the highly integrated networks operating between different pluripotency factors, about 6% and 11% of Sox2 interactors from this study were also a part of protein complexes found interacting with Nanog (Wang et al., 2006) and Oct4 (van den Berg et al., 2010) respectively (Supplementary Table 7). Therefore, most of the associations reported here are novel. Our results highlight putative novel functions of the transcription factor Sox2 as a constituent of ribonucleoprotein complexes containing RNA splicing and processing proteins, which is in line with the increasing connection between transcriptional regulation and RNA processing factors (Pandit et al., 2008)

Our data also indicated that Sox2 in present in complexed which include specific RNAs, such as mRNAs and the ncRNA 7SK. We have previously shown that 7SK represses a subset of genes with active or bivalent chromatin marks in mES cells, along with those involved in lineage specification (Castelo-Branco et al., 2013). Both Sox2 and Sox10 have been shown to regulate transcriptional elongation of myelin genes in Schwann cells by interacting directly with P-TEFb (Arter and Wegner, 2015), which is a primary regulatory target of 7SK. In addition, Poly ADP-ribose polymerase 1 (PARP-1), another Sox2 interactor in our study, was recently shown to facilitate and stabilize Sox2 binding to high nucleosome harbouring euchromatic regions (Liu and Kraus, 2017). PARP-1 also ADP-ribosylates and inhibits the negative elongation factor (NELF), thereby allowing transcriptional elongation to proceed (Gibson et al., 2016). Previous studies have also hinted at KAP1/Trim28 (interactor of Sox2 in this study) mediated recruitment of inactive P-TEFb in complex with 7SK to promoter proximal regions needing a transcription factor or other DNA binding proteins to interface with chromatin (D’Orso, 2016). Hence, the association of 7SK with Sox2 could be similarly important in modulating transcriptional programs dependent on Sox2 in ES cells. ATAC-seq following knock down of 7SK in mouse ES cells resulted in a reduction of Sox2 transcription factor footprint on enhancer elements (Flynn et al., 2016). Nevertheless, our data indicates that such a function would not be dependent on mutual modulation of recruitment to chromatin.

Long non-coding RNAs are now thought to be integral to the pluripotency circuit of ES cells (Dinger et al., 2008; Guttman et al., 2011; Loewer et al., 2010). LncRNAs involved in pluripotency maintenance and neurogenesis (Ng et al., 2012) including lncRNA RMST were shown to interact with Sox2 (Ng et al., 2013). Previous studies investigating Sox2 protein interactome in ES cells as well as other cell types have also found proteins with RNA binding capability (Cox et al., 2013; Fang et al., 2011; Gao et al., 2012; Zhou et al., 2016) with one in-vitro study implicating the Sox2 HMG domain in binding RNA (Tung et al., 2010). We detect a limited number of RNAs interacting with Sox2, which includes ncRNAs, 7SK and Snord34. Our interactome analysis indicates two RNA-binding proteins that could mediate association of Sox2 with 7SK, namely Srsf1 and hnRNAPA2/B1. Srsf1 along with Srsf2, were shown to associate with gene promoters in a 7SK dependent manner and play a direct role in transcription pause release (Ji et al., 2013). HnRNPA2/B1 specifically interacts in the nucleoplasm with a portion of 7SK that is not in complex with its canonical partners, HEXIM1 and P-TEFb, and is involved in dynamic remodeling of 7SK snRNP (Barrandon et al., 2007; Van Herreweghe et al., 2007). Thus Sox2 might be involved in processes downstream of transcriptional initiation. It is also possible that interaction of Sox2 with snoRNAs and mRNAs might regulate other chromatin related processes. Interestingly, snoRNAs have been recently shown to be present at the chromatin (Li, Zhou et al., 2017, Sridhar, Rivas-Astroza et al., 2017) and regulate chromatin/nuclear structure (Schubert, Pusch et al., 2012). Alternatively, it remains a possibility that the association between Sox2 and the RNAs reported here is a consequence of their proximity on DNA and nucleoplasm and not necessarily due to any functional relationship. Future investigations might unveil whether the presence of Sox2 in ribonucleoprotein complex carries any significance either to the functionality of Sox2 or its partner RNAs.

Our results indicate that Sox2 is associated with several complexes in the chromatin and nucleoplasm in mouse ES cells, including ribonucleic complexes. While our data suggests that the interaction of Sox2 with the ncRNA 7SK does not regulate their recruitment to chromatin, it is possible that this crosstalk represents a new facet for the mechanism of action of Sox2 in the nucleoplasm and at the chromatin.

## Methods

### Cell culture

J1 expressing the biotin ligase BirA and Sox2, Nanog or Oct4 flanked by a peptide amenable to biotinylation by BirA, as well as control J1 cells only expressing BirA, were kindly provided by Dr. Stuart H. Orkin (Dana Farber, Harvard Medical School) (Kim et al., 2009; Wang et al., 2006). Briefly, this in-vivo biotinylation system was set up with a J1 mES cell line stably expressing the bacterial BirA gene. BirA-expressing cells were subsequently used to introduce a plasmid encoding a peptide-substrate for the BirA enzyme fused to the transcription factor of interest, to produce stably expressing biotinylated transcription factor (bioTF) mES cell lines.

2TS22C mES cells, where Sox2 can be deleted upon doxycycline treatment, were kindly provided by Dr. Hitoshi Niwa at the RIKEN Center for Developmental Biology, Kobe, Japan (Masui et al., 2007). All mES cell lines were grown on 0.1% gelatin coated plates and maintained in ES media consisting of Glasgow Minimum Essential Medium (GMEM) supplemented with 10% fetal calf serum for ESCs (Biosera, Boussen, France), 0.1 mmol/l nonessential amino acids, 2 mmol/l L-Glutamine, 1 mmol/l sodium pyruvate, 0.1 mmol/l β-mercaptoethanol, 1x penicillin/streptomycin and 10^6^ units/l LIF (ESGRO, MilliporeCorp., Billerica, MA, USA). For SILAC experiments, SILAC Advanced DMEM/F12 media was used (Invitrogen, SILAC Protein ID and Quantification Kit, MS10033). For Sox2 deletion, 2TS22C mES cells were treated with 1 μg/ml doxycycline for 24 h.

### SILAC quantitative proteomics

BioSox2 expressing J1 ES cells along control cells were grown in either light (^12^C_6_) or heavy medium (^13^C_6_) for 6 passages. The cells were collected by accutase treatment and washed twice with ice-cold PBS. The pellet was resuspended in 5 packed cell volumes (pcv) of ice-cold nuclear extract buffer A without NP-40 (all buffer compositions are included in Supplementary word file 1). After spinning for 10 min at 2,400 g at 4°C, the pellet was resuspended in 3 pcv of ice-cold nuclear extract buffer A with NP-40. After incubating the cells at 4°C with gentle rotation, they were homogenized with 10 strokes of Dounce homogenizer (type B, wheaton 1 ml). Nuclei were pelleted by centrifugation at 4°C for 15 min at 4,300 g. The resulting supernatant (cytoplasmic fraction) was removed and the pellet was resuspended in 2 nuclear pellet volumes (npv) of ice-cold nuclear extract buffer B followed by homogenization and extraction of nuclei for 1 h at 4°C with gentle rotation. After centrifugation at 13,200 rpm for 30 min, the supernatant (nuclear extract) was transferred to a new tube and the pellet (chromatin) was resuspended in 350 μl digestion buffer (Active Motif, ChIP-IT Enzymatic Kit, catalogue number 53006) supplemented with 7.9 μl PIC, 7.9 μl PMSF and 0.875 μl SuperaseIN RNAse inhibitor (ThermoFisher Scientific, AM2696). Chromatin samples were incubated for 5 min at 37°C followed by a second incubation for 10 min at 37°C with shaking at 1,000 rpm after the addition of 1:100 enzymatic working solution (Active Motif, ChIP-IT Enzymatic Kit, catalogue number 53006). The reaction was stopped with the addition of 7 μl EDTA 0.5 M and the samples were chilled on ice for 10 min. Supernatant was collected after centrifugation at 12,000 rpm (4°C) for 12 min and protein concentration was measured. Equal amounts of protein from chromatin fractions of control and bioSox2 were used for IP. The nuclear extract was ultracentrifuged for 1hr at 60,000 g at 4°C. Supernatant was collected, protein concentration was measured and equal amounts of protein from nuclear fractions of control and bioSox2 were used for IP.

50 μl of Protein G dynabeads (per 5 mg protein) was washed with ice cold nuclear extract buffer B (nuclear extract) or digestion buffer (chromatin), resuspended in respective buffers and 50 μl was used to pre-clear the extracts for 1hr at 4°C with gentle rotation. 50 μl of Dynabeads MyOne Streptavidin T1 (ThermoFisher Scientific) was washed and resuspended as previously indicated and 50 μl was added to the pre-cleared supernatant and incubated overnight at 4°C with gentle rotation. The beads were washed twice with IP350 0.3 % buffer for 15 min with gentle rotation at 4°C, beads from control and bioSox2 were mixed before the final wash for both chromatin and nuclear fractions, then were eluted in 2x SDS sample buffer. This was followed by heating at 95°C for 5 min, vortexing, cooling to RT and pelleting the beads. The elution was repeated with 1xSDS sample buffer. Supernatants were pooled and the beads were pelleted into 4xNuPAGE loading buffer. Extracted proteins were resuspended in Laemmli Sample Buffer, and resolved on a 4-20 % SDS-PAGE. The gel was stained with Coomassie blue, cut into 20 slices and processed for mass spectrometric analysis using standard in gel procedure. Briefly, cysteines were reduced with dithiothreitol (DTT), alkylated using chloroacetamide (CAA) (Nielsen et al., 2008), and finally the proteins were digested overnight with endoproteinase Lys-C and loaded onto C18 StageTips prior to mass spectrometric analysis.

### LC/MS

All MS experiments were performed on a nanoscale EASY-nLC 1000 UHPLC system (Thermo Fisher Scientific) connected to an Orbitrap Q-Exactive Plus equipped with a nanoelectrospray source (Thermo Fisher Scientific). Each peptide fraction was eluted off the StageTip, auto-sampled and separated on a 15 cm analytical column (75 μm inner diameter) in-house packed with 1.9 μm C18 beads (Reprosil Pur-AQ, Dr. Maisch) using a 75 min gradient ranging from 5 % to 40 % acetonitrile in 0.5 % formic acid at a flow rate of 250 nl/min. The effluent from the HPLC was directly electrosprayed into the mass spectrometer. The Q Exactive plus mass spectrometer was operated in data-dependent acquisition mode and all samples were analyzed using previously described ‘sensitive’ acquisition method (Kelstrup et al., 2012). Back-bone fragmentation of eluting peptide species were obtained using higher-energy collisional dissociation (HCD) which ensured high-mass accuracy on both precursor and fragment ions.

### Identification of peptides and proteins by MaxQuant

The data analysis was performed with the MaxQuant software suite (version 1.3.0.5) as described (Cox and Mann, 2008) supported by Andromeda (www.maxquant.org) as the database search engine for peptide identifications (Weidner et al., 1990). We followed the step-by-step protocol of the MaxQuant software suite (Cox et al., 2009) to generate MS/MS peak lists that were filtered to contain at most six peaks per 100 Da interval and searched by Andromeda against a concatenated target/decoy (forward and reversed) version of the IPI human database. Protein sequences of common contaminants such as human keratins and proteases used were added to the database. The initial mass tolerance in MS mode was set to 7 ppm and MS/MS mass tolerance was set to 20 ppm. To minimize false identifications, all top-scoring peptide assignments made by Mascot were filtered based on previous knowledge of individual peptide mass error. Peptide assignments were statistically evaluated in a Bayesian model on the basis of sequence length and Andromeda score. We only accepted peptides and proteins with a false discovery rate of less than 1 %, estimated on the basis of the number of accepted reverse hits.

### Gene ontology analysis

Candidates that showed at least two-fold enrichment over control in the forward and reverse labelling in SILAC experiments were considered for analysis. GO analysis was performed with DAVID 6.7 (Huang et al., 2009). *P*-values were adjusted for multiple hypothesis testing using the Benjamini-Hochberg method. Significantly enriched categories in the subontology of functional category, pathways and protein domains with an adjusted *P*-value < 0.05 were chosen.

### Co-immunoprecipitation of GST tagged HP1 proteins with recombinant Sox2 or ES cell nuclear extract

Recombinant proteins were expressed in and purified from *Escherichia coli* as described previously (Bannister and Kouzarides, 1996). Mouse full-length HP1 isoforms and the chromo domain (residues 5–80), hinge (residues 61–121) and chromo-shadow domain (residues 110–188) of HP1a were cloned into pGex vector and expressed as a GST fusion protein. Glutathione sepharose beads were prepared by washing 1 ml of beads (5 mg GST capacity) with 5 ml GST buffer, spinning at for 5 min at 500 g at 4°C and resuspending in 1 ml GST buffer (50 % slurry, Vf = 2 ml, capacity 2.5 μg/μl). 20 μl 50 % slurry glutathione sepharose beads was added to low binding tubes, together with 485 μl GST buffer, 0.5 μg recombinant human Sox2 (Abcam ab95847), and 5-10 μg GST, 10 μg GST-HP1α, 10 μg GST-HP1β or 10 μg GST-HP1γ. Alternatively, 147.5 μl GST buffer was added to low binding tubes, together with 0.5 μg recombinant Sox2 (Abcam ab95847), and 5-10 μg GST + <u>2 μl</u> 50 % slurry glutathione sepharose beads, and 10 μg of GST-HP1α-FL, GST-HP1α-CSD, GST-HP1α-CD, GST-HP1α-H, GST-HP1β-FL, GST-HP1β-CSD, GST-HP1β-CD or GST-HP1β-H, in glutathione sepharose beads. Samples were incubated for 2 h at 4°C with end-to-end rotation, spinned for 5 min at 500 g at 4°C. Beads were washed four times with 1 ml GST lysis buffer (with spins for 5 min at 500 g at 4°C). GST fusion and bound proteins were eluted with 30 μl 2xLaemmli buffer and boiled for 5 min, prior to western blot.

For co-IPs with mouse ES cell nuclear extracts (isolated as described in the quantitative proteomics section), these were pre-cleared and RNAse treated by incubating 25 μg GST protein, 20 μl 50 % slurry glutathione sepharose beads (50 μg capacity), 200 μg Oct4 GIP ES nuclear extracts, 5 μl RNase A (2.5 g, DNase-free, Roche #11119915001) or dH_2_O, and GST buffer. Samples were incubated for 1 h at room temperature and centrifuged for 5 min at 500 g at 4°C. The pre-cleared supernatants were then mixed with 20 μl 50 % slurry glutathione sepharose beads, and 5-10 μg GST, 10 μg GST-HP1α, 10 μg GST-HP1β or 10 μg GST-HP1γ in glutathione sepharose beads. Samples were incubated for 2 h at 4°C and centrifuged for 5 min at 500 g at 4°C. Beads were washed twice with 1 ml GST lysis buffer and twice with 0.5 ml GST lysis buffer (with spins for 5 min at 500 g at 4°C). GST fusion and bound proteins were eluted with 30 μl 2xLaemmli buffer and boiled for 5 min, prior to western blot.

### Western blot

Cell monolayers or pellets were resuspended in 2xLaemmli buffer, boiled for 5 min at 95°C and passed 10 times through a 21 G needle to shear genomic DNA. Proteins were separated by SDS–PAGE, transferred to nitrocellulose membrane (Millipore) using wet transfer and incubated in blocking solution (5 % BSA in TBS containing 0.1 % Tween) for 1hr at room temperature. Membranes were incubated with primary antibody at 4°C overnight and appropriate HRP-conjugated secondary antibody for 2 h at room temperature. Membranes were then incubated for chemiluminescence (ECLH; GE Healthcare) and proteins were detected by exposure to X-ray film. Primary antibodies, diluted in blocking solution were used against Sox2 (α-Sox2, raised in goat, Y-17, Santa Cruz, sc-17320).

### RNA immunoprecipitation (RIP) and sequencing

All incubations were performed in low-bind RNase-free tubes. 50 million cells/IP were fixed with 1 % formaldehyde (Sigma F8775) for 10 min at room temperature, quenched with Glycine stop solution (Active Motif, ChIP-IT Enzymatic Kit, catalogue number 53006) and lysed. The total nuclear lysate was either sonicated at high frequency (H 20W) with the 30 s ON/30 s OFF setting for 20 min in BioRuptor or processed for nuclei isolation and enzymatic digestion, as described for the SILAC quantitative proteomics (Active Motif, ChIP-IT Enzymatic Kit, catalogue number 53006). Sheared chromatin was pre-cleared with Protein G Dynabeads for 1 h at 4°C with gentle rotation. Immunoprecipitation of biotinylated Sox2/Oct4/Nanog was performed with 50 μl MyOne streptavidin Dynabeads T1 overnight followed by washing with FA1000, LiCl and TES buffers. RNA was eluted after reverse-crosslinking (65°C for 1 h with 1,000 rpm rotation), Qiazol was added and RNA was extracted using the Qiagen miRNAeasy kit. Sample preparation for sequencing was done by either adapting the directional mRNA-Seq protocol (Illumina RS-100-0801) to the small RNA Illumina sequencing, v1.5 small RNA 3’ adaptor kit (Illumina FC-102-1009) or by using TruSeq directional small RNA kit (Illumina RS-200-0012). In order to capture both long and short RNAs, RNA was fragmented (Ambion AM8740) prior to sample preparation. Total RNA was depleted from ribosomal RNA by treatment with Ribo-Zero rRNA removal kit (RZH1086). Sequencing was performed with Illumina instruments to obtain single-end or paired-end reads (Supplementary Table 3). For endogenous RIPs, the following antibodies were used: mouse∝-FLAG M2 (Sigma), Sox2 (α-Sox2, raised in goat, Y-17, Santa Cruz, sc-17320), Nanog Antibody AbVantage Pack (Bethyl, A310-110A), Mouse KLF4 Affinity Purified Polyclonal Ab, Goat IgG (AF3158, R&D Systems), Oct-3/4 (N-19) X, Polyclonal Antibody (sc-8628-X, Santa Cruz) and Suz12 (Abcam, ab12073).

### RNA-seq data processing and analysis

RNA-seq data from RIP and input samples were processed in the same manner, using the best-practice RNA-seq pipeline from the National Genomics Infrastructure Sweden (NGI-RNAseq v1.4; https://github.com/SciLifeLab/NGI-RNAseq), including adapter trimming with cutadapt v1.16 (Martin, 2011), mapping to mouse genome assembly GRCm38 with STAR v2.5.3a (Dobin et al. 2013), counting reads per gene (Ensembl release 92 annotation) with featureCounts v1.6.0 (Liao et al., 2014), and multiple quality control steps. Read counts were normalized among samples using the size factor method implemented in the BioConductor package DESeq2 (Anders and Huber, 2010). To identify differences in gene expression between bioSox2 and control J1 cells, the input samples were compared using DESeq2 v1.22.2 with default parameters, including experimental batch as a factor to account for differences in library preparation and sequencing between the two batches. P-values were adjusted by the Benjamini-Hochberg method to control the false discovery rate (FDR). To identify RNAs enriched by RIP, an enrichment ratio was computed per batch, as (bioSox2 RIP / control RIP) / (bioSox2 input / control input), using normalized counts incremented by a pseudo-count of 0.1 to avoid denominators of zero. RNAs with enrichment ratio > 2 and bioSox2 RIP raw read count > 50 in both batches were considered hits.

### qRT-PCR

Total RNA was extracted using the miRNeasy Extraction Kit (Qiagen), with in-column DNAse treatment. 500 ng of RNA was reverse transcribed using the High capacity cDNA reverse transcription kit (4368814, Applied Biosystems) including RNase inhibitor (N8080119, Applied Biosystems). A reverse transcriptase negative (RT-) control was included for each sample. Both the cDNA and the RT-were diluted 1:3 in RNase/DNAse free water for qRT-PCR. qRT-PCR reactions were run on a StepOnePlus™ System (Applied Biosystems) in duplicate and with RT-reactions to control for genomic DNA. Fast SYBR^®^ Green Master Mix (4385616, Applied Biosystems) was used according to the manufacturer’s instructions; each PCR reaction had a final volume of 10 μl with 2.5 μl of diluted cDNA or RT-. The running conditions were 20 s at 95°C, followed by 40 cycles of 3 s of 95°C and 30 s of 60°C, then 15 s at 95°C, 1 min at 60°C and 15 s at 95°C. *Tbp* was run as housekeeping gene. Double delta Ct method was used for calculating fold change.

### Chromatin Isolation by RNA Purification (ChIRP)

ChIRP was performed as previously described (Chu et al., 2012). Mouse 2TS22C cells were cultured as above and either treated with Dimethyl sulfoxide (DMSO) or Doxycycline (1 μg/ml) for 24 h before cross-linking with glutaraldehyde. 20 million cells were used per ChIRP. Six probes covering the whole length of 7SK were used and depending on their positions along the RNA were divided into odd and even probe pools (Flynn et al., 2016). A single probe against LacZ mRNA was used as a negative control. Isolated RNA from a small aliquot of post-ChIRP beads was used in qRT-PCR to quantify 7SK enrichment. Isolated DNA following ChIRP was used to make sequencing library with ThruPLEX DNA-seq 12S kit (R400428, Rubicon Genomics). The library was quantified with KAPA library quantification kit (Illumina), samples were pooled and then sequenced on HiSeq2500 at National Genomics Infrastructure (NGI), SciLife Lab, Stockholm.

### ChIRP-seq data analysis

Sequence reads were trimmed using trim_galore v0.4.0 (Krueger, 2012) to remove adapter sequences and low quality bases from the 3’ end of the reads. Reads less than 20 bp were removed post-trimming, prior to mapping. Trimmed reads were mapped to the mm10 mouse genome from the UCSC database using bwa v0.7.12 (Li and Durbin, 2009) with parameters -n 3 -k 2 -R 300. Peak calling was performed for each ChIRP pulldown using macs2 (Zhang et al., 2008) with parameter -q 0.001 using the corresponding Input sample as control. Downstream analyses were conducted using the Bioconductor suite of packages (Huber et al., 2015) in R (R core team, 2017). Robust 7SK binding sites were identified by taking the overlap between the peaks called using the odd and even probe pools. Peaks that also overlapped a peak from the LacZ negative control were removed. A final set of 7SK binding sites was identified by taking the union between the doxycycline treated and untreated filtered probe sets. Annotation of our peaks and those from external data sets was performed against the UCSC mm10 knownGene database using the clusterProfiler package (Yu et al., 2012). Target genes were identified based on overlap of significant peaks with either the gene body or the promoter region defined as the region 2.5 Kb upstream of the TSS. Quantification of ChIRP signal at loci of interest was performed using modified scripts from the Repitools package (Statham et al., 2010).

### Chromatin Immunoprecipitation (ChIP)

Briefly 300,000 2TS22C cells were plated per condition in a 6-well plate. 100 nM of scrambled ASO or 7SK 3’ ASO (IDT) were transfected using Lipofectamine 2000 (Invitrogen) using the manufacturer’s recommendations. Opti-MEM reduced serum medium was used to prepare the complexes. Cells were incubated with these complexes overnight before replacing with fresh medium. After 24 h, cells were either collected into Qiazol (Qiagen) for RNA extraction or were cross-linked with 1 % formaldehyde (37 %, Sigma-Aldrich) for ChIP.

The protocol and buffers from the True MicroChIP kit (C01010130, Diagenode) were used to perform sonication and immunoprecipitation (IP) with 100,000 cells per condition. Cells were sheared for 25 min using Bioruptor (Diagenode) with 30s ON/30s OFF setting under high power (H). 0.5 μg of Sox2 antibody (AF2018, R&D) or goat IgG was used for each IP. The immune complexes were purified with DiaMag Protein G coated magnetic beads (C03010021, Diagenode). De-crosslinked DNA was eluted for qPCR to assess changes in Sox2 recruitment to specific areas of interest following 7SK knock down. To compare Sox2 recruitment between control and 7SK depleted cells, the qPCR data was normalized to 10 % purified input DNA, which was used as a measure of total chromatin present in the particular sample.

## Supporting information

List of Buffers

## Acknowledgements

We would like to thank Alessandra Nanni, Tony Jimenez-Beristain, Ahmad Moshref and Johnny Söderlund for support, Mark A. Dawnson and Andy Bannister for providing HP1 recombinant proteins, Daniel Gaffney (Wellcome Sanger Institute) and José Silva (Wellcome-MRC Cambridge Stem Cell Institute) for discussions. We thank the National Genomics Infrastructure and Uppmax for providing assistance in massive parallel sequencing and computational infrastructure. The bioinformatics computations were performed on resources provided by the Swedish National Infrastructure for Computing (SNIC) at UPPMAX, Uppsala University. Work in G.C.-B.’s research group was supported by Swedish Research Council (grants 2010-3114), European Union (FP7/Marie Curie Integration Grant EPIOPC, Horizon 2020 Research and Innovation Programme/European Research Council Consolidator Grant EPIScOPE, grant agreement number 681893), Ming Wai Lau Centre for Reparative Medicine, and Karolinska Institutet. The sequencing data for both RIP-seq and ChIRP-seq are deposited in ArrayExpress bearing accession numbers E-MTAB-7640 and E-MTAB-7570, respectively.

## Author Contributions

G.C.B, P.A and T.K. conceptualized the study. G.C.B performed the SILAC-MS, RIP, co-immunoprecipitation experiments, S. performed ChIRP and ChIP experiments, P.A. co-performed some of the previous and additional experiments. M.L.N. analysed the MS data. P.E and S.R performed bioinformatics analyses on sequencing data from RIP and ChIRP. G.C.B and S wrote the paper with input from all the authors.

---

**Supplementary figure 1**

A) Gene expression correlation between control and bioSox2 mES J1 cell lines measured by RNA-seq. Normalized read counts are plotted for all detected genes, comparing the control and bioSox2 input samples (mean across the two experiments). Red circles indicate differentially expressed genes (FDR-adjusted *P* < 0.1), listed in Supplementary Table 1.

B) Mass-spectrometric chromatogram of HP1 peptide showing peaks from LIGHT amino acid labelled control (red) and HEAVY amino acid labelled bioSox2 (blue) chromatin extracts.

C) Western blot indicating pull down of Sox2 in co-immunoprecipitation experiments with GST-tagged HP1 α, β, γ proteins and recombinant human Sox2, compared to the control. (n=2 for HP1α)

D) Representative western blot indicating pull down of Sox2 from ES nuclear extract in immunoprecipitation experiments with GST-tagged HP1 α, β, γ proteins in the presence or absence of RNase (n=2 for HP1α and β)

E) Western blot indicating successful pull down of Sox2 in co-immunoprecipitation experiments with different GST-tagged domains of HP1 α, β proteins (FL-full length, CSD-chromo shadow domain, CD-chromo domain) and recombinant human Sox2. Different domains of HP1 proteins exhibit varying affinities for Sox2 (n=1)

F) RT-qPCR showing enrichment of *7SK* and *Snord34* RNAs pulled down in ES cell following (left) RNA immunoprecipitation with biotin tagged Sox2 and other pluripotency factors, bioOct4 and bioNanog; Y-axis, % of input (right); RNA immunoprecipitation of endogenous proteins with antibodies against Sox2, Oct4, Nanog, Klf4 and Suz12. Y-axis, fold enrichment to FLAG IP.

G) RT-qPCR showing enrichment of *Sox2, Pou5f1* (Oct4), *Nanog* and *Suz12* mRNAs pulled down in ES cell following (above) RNA immunoprecipitation with biotin tagged Sox2 and other pluripotency factors, bioOct4 and bioNanog; Y-axis, % of input (below); RNA immunoprecipitation of endogenous proteins with antibodies against Sox2, Oct4, Nanog, Klf4 and Suz12. Y-axis, fold enrichment to FLAG IP.

