## Supplementary material for "Interaction of Sox2 with RNA binding proteins in mouse embryonic stem cells": List of Buffers

**Biotin IP (SILAC)**

Nuclear Extract A (hypotonic buffer) (at 4°C):

**20mM HEPES pH 7.5**

**1mM EDTA**

**10mM KCl**

**0,2% Nonidet P-40 v/v**

**10% glycerol (v/v)**

**1mM DTT**

**1mM PMSF**

**1x Complete Protease inhibitors**

**50U/ml SuperASin (Ambion)**

**ddH_2_O**

Nuclear Extract B (at 4°C):

**20mM HEPES pH 7.5^[1]^**

**350mM NaCl**

**1mM EDTA**

**10mM KCl**

**20% glycerol (v/v)**

**1mM DTT**

**1mM PMSF**

**1x Complete Protease inhibitors**

**50U/ml SuperASin (Ambion)**

**ddH_2_O**

IP 350 buffer (at 4°C):

**20mM Tris-HCl pH 7.5**

**350mM NaCl**

**1mM EDTA**

**0,3% Nonidet P-40 v/v**

**0,5% Nonidet P-40 v/v**

**10% glycerol (v/v)**

**1mM DTT**

**0,2mM PMSF**

**1x Complete Protease inhibitors**

**50U/ml SuperASin (Ambion)**

**ddH_2_O (0,3)**

**ddH_2_O (0,5)**

2x SDS sample buffer:

**0.5M Tris-HCl pH 6.8**

**3% SDS v/v**

**1% glycerol (v/v)**

**ddH_2_O**

**GST coIP**

Dialysis Buffer:

**20mM HEPES (KOH) pH 7,5**

**20% glycerol**

**100mM KCl**

**0,83mM EDTA**

**1,66mM DTT**

**0,2mM PMSF**

**DEPC ddH_2_O**

GST Buffer (at 4°C):

**25mM HEPES pH 7.5^[2]^**

**150mM NaCl**

**0,1% Nonidet P-40 v/v**

**1mM EDTA**

**10% glycerol (v/v)**

**1mM PMSF**

**1x Complete Protease inhibitors**

**ddH_2_O**

**RNA Immunoprecipitation (RIP)**

Lysis buffer (at 4°C):

**50mM HEPES pH 7.5**

**1mM EDTA**

**1% Triton X-100**

**140mM NaCl**

**0,1% NaDeoxycholate**

**1x Complete Protease inhibitors EDTA- free**

**50U/ml SuperASin (Ambion)**

**DEPC ddH_2_O**

Binding buffer (at 4°C):

**50mM HEPES pH 7.5**

**20mM EDTA**

**0,5% Triton X**

**25mM MgCl_2_**

**5mM CaCl_2_**

**50U/ml SuperASin (Ambion)**

**DEPC ddH_2_O**

FA1000 buffer (at 4°C):

**50mM HEPES pH 7.5**

**1mM EDTA**

**1% Triton X**

**1M NaCl**

**0,1% NaDeoxycholate**

**50U/ml SuperASin (Ambion)**

**DEPC ddH_2_O**

LiCl buffer (at 4°C):

**10mM Tris-HCl pH 7.5**

**1mM EDTA**

**1% Triton X**

**250mM LiCl**

**0,5% NaDeoxycholate**

**50U/ml SuperASin (Ambion)**

**DEPC ddH_2_O**

TES buffer (at 4°C):

**10mM Tris-HCl pH 7.5**

**1mM EDTA**

**10mM NaCl**

**50U/ml SuperASin (Ambion)**

**DEPC ddH_2_O**

Elution buffer:

**50mM Tris-HCl pH 8.1**

**10mM EDTA pH8.0**

**1% SDS**

**50U/ml SuperASin (Ambion)**

**DEPC ddH_2_O**
