## Supplementary figures and images for "Interaction of Sox2 with RNA binding proteins in mouse embryonic stem cells"

### Supplementary Figure 1

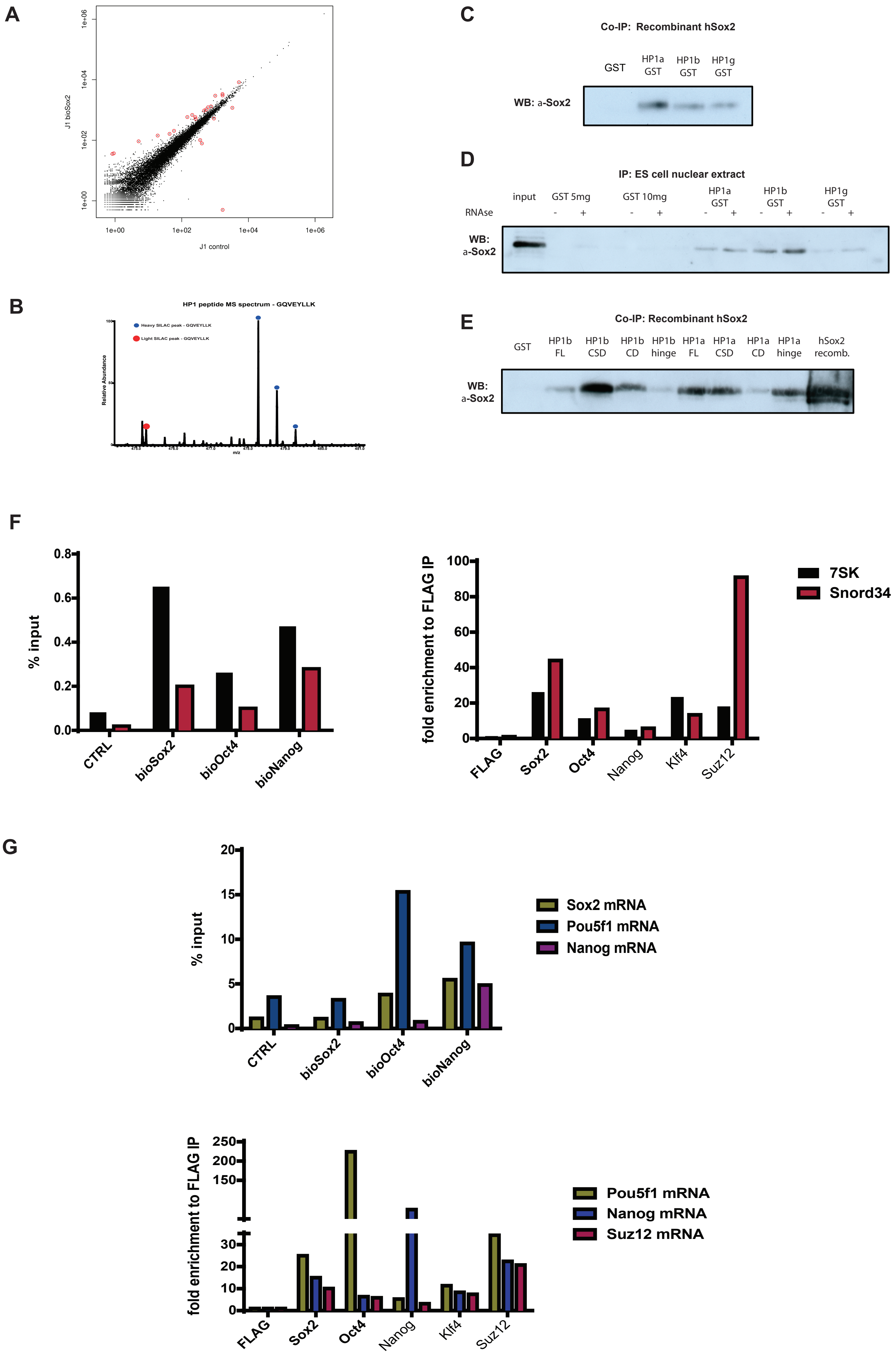
